# Characterization of a novel estrogen- and progesterone-responsive endometrial cancer cell line: HCI-EC-23

**DOI:** 10.1101/2022.08.25.505203

**Authors:** Craig M. Rush, Zannel Blanchard, Jacob T. Polaski, Kyle S. Osborne, Krystle Osby, Jeffery M. Vahrenkamp, Chieh-Hsiang Yang, David H. Lum, Christy R. Hagan, Kimberly K. Leslie, Miles A. Pufall, Kristina W. Thiel, Jason Gertz

## Abstract

Most endometrial cancers express the hormone receptor estrogen receptor alpha (ER) and are driven by excess estrogen signaling. However, evaluation of the estrogen response in endometrial cancer cells has been limited by the availability of hormonally responsive *in vitro* models, with one cell line, Ishikawa, being used in most studies. Here, we describe a novel, adherent endometrioid endometrial cancer (EEC) cell line model, HCI-EC-23. We show that HCI-EC-23 retains ER expression and that ER functionally responds to estrogen induction over a range of passages. We also demonstrate that this cell line retains paradoxical activation of ER by tamoxifen, which is also observed in Ishikawa and is consistent with clinical data. The mutational landscape shows that HCI-EC-23 is mutated at many of the commonly altered genes in EEC, has relatively few copy-number alterations, and is microsatellite instable high (MSI-high). *In vitro* proliferation of HCI-EC-23 is strongly reduced upon combination estrogen and progesterone treatment. HCI-EC-23 exhibits strong estrogen dependence for tumor growth *in vivo* and tumor size is reduced by combination estrogen and progesterone treatment. Molecular characterization of estrogen induction in HCI-EC-23 revealed hundreds of estrogen-responsive genes that significantly overlapped with those regulated in Ishikawa. Analysis of ER genome binding identified similar patterns in HCI-EC-23 and Ishikawa, although ER exhibited more bound sites in Ishikawa. This study demonstrates that HCI-EC-23 is an estrogen- and progesterone-responsive cell line model that can be used to study the hormonal aspects of endometrial cancer.

## Introduction

Cancer of the uterine corpus is the most common gynecologic cancer, 4th most common overall cancer type, and 6th most common cause of cancer death for women in the US, with an expected 65,950 new cases and 12,550 deaths in 2022[1]. Uterine cancer remains one of the few cancers with increasing incidence and mortality, in part due to its strong association with the obesity epidemic[2,3]. The vast majority (>90%) of cancers of the uterine corpus are cancers of the endometrium, the inner epithelial lining of the uterus. Endometrial cancers are broadly separated into two histopathological subtypes[4]. Type I tumors make up ~85% of endometrial tumors and have low-grade (grade I or II) endometrioid histology. Type I tumors express high levels of estrogen receptor alpha (ER) and are believed to be hormonally driven with excess estrogen signaling being a significant risk factor. In contrast, type II tumors are thought to be hormone-independent and include high-grade (grade III) endometrioid tumors along with tumors of other histologic subtypes, including serous, clear cell, carcinosarcoma, and mixed histology. The broad histopathological classification of endometrial cancer also associates with molecular classifications identified by The Cancer Genome Atlas, where type I tumors fall largely into the copy-number low, microsatellite instable (MSI), and *POLE* mutated etiologies, while type II tumors are enriched for the copy-number high etiology and are enriched for *TP53* mutations[5–7].

There have been many cell lines established from type I tumors in order to model disease biology, such as Ishikawa[8,9], AN3CA[10], HEC-1-A[11], RL95-2[12], MFE-280[13], MFE-296[13], JHUAS-1[14], and JHUEM-2[14]. However, the majority of these lines have limited ability to assess the estrogen driven aspects of type I endometrial cancer biology due to loss of ER expression and response to estrogens. This has left the Ishikawa cell line as the prevailing model for studying the hormonal aspects of endometrial cancer *in vitro*[15–22]. Early studies made use of another hormone receptor positive cell line, ECC-1, which has been recently identified to be contaminated by the Ishikawa line at some point, rendering its utility limited[23,24]. The lack of additional cell lines to study the >85% of endometrial cancers that are hormonally-driven has limited the generalizability of previous studies. This is especially true when considering the example of hormone receptor positive breast cancer cell lines, where there are variable responses to hormones, therapeutic responses, and resistance mechanisms to anti-hormone therapy[25–29]. Many of these differences can be attributed to the unique genetic background of each of the cell lines[29]. Therefore, we are limited to studying the estrogen-driven aspects of type I endometrial cancer through the lens of a single genetic background, that of the MSI-high Ishikawa cell line[24]. The increasing use of patient-derived xenograft (PDX) models in endometrial cancer has increased the availability of hormone-responsive model systems[30–33]. Although these PDX models are highly informative, cell lines remain the predominant model system where large numbers of cells are needed, such as drug and genetic screens, or for genetic manipulation to model clinically-relevant mutations.

We addressed the lack of hormonally-responsive endometrial cancer cell lines by isolating and characterizing endometrial cancer cell lines from PDX models, which led to the development of a cell line model, HCI-EC-23, that retains estrogen-responsiveness *in vitro* and *in vivo*. We demonstrate that this cell line is epithelial in nature and grows well as a monolayer *in vitro*. We also describe the genetic background of HCI-EC-23 by showing that it is mutated at many of the top mutated genes in endometrioid endometrial carcinoma and that it is MSI-high. Critically, the HCI-EC-23 cell line retains ER expression and estrogen-responsiveness *in vitro* and *in vivo* as assayed by western blot, luciferase reporter assays, growth assays, and RNA-seq. We also show that the HCI-EC-23 is similar but distinct from Ishikawa in its response to 17β-estradiol (E2) on a transcriptional level and where ER binds to the genome. We show that ER in both HCI-EC-23 and Ishikawa is paradoxically activated by tamoxifen, while HCI-EC-23 shows an anti-proliferative response to progesterone (P4). This study details the characterization of a novel endometrial cancer cell line that retains hormone-responsiveness and will advance the field’s understanding of the hormonal underpinnings of endometrial cancer by providing an additional yet novel model system.

## Results

### Generation of PDX-derived cell lines

Endometrioid endometrial carcinoma (EEC) is commonly believed to be driven by excess estrogen signaling. Despite the fact that most EEC cell models are derived from tumors expressing ER[8–14], most of these lines have lost ER expression at the protein level (**Figure 1a**). The only commercial line that has retained estrogen responsiveness, allowing for modeling of the hormonal action of endometrial cancer, is Ishikawa **(Figure 1a)**[15–22].

**Figure 1.**
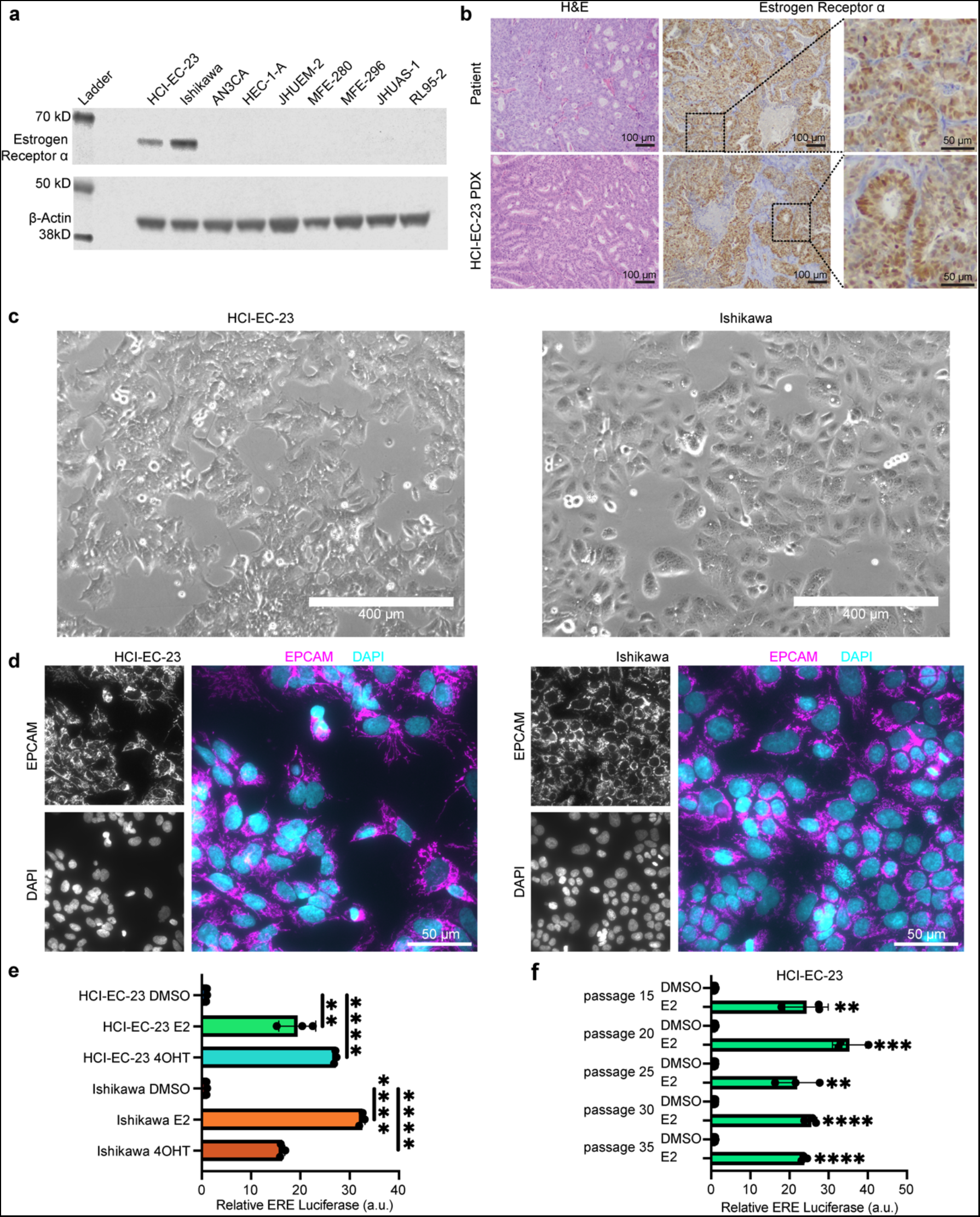
HCI-EC-23 is an epithelial endometrial cancer cell line expressing ER. **(a)** Western blot shows protein levels of ER and loading control β-actin. Cell lines shown are HCI-EC-23 along with commonly used, commercially available endometrial cancer cell lines. The only cell lines expressing ER are HCI-EC-23 and Ishikawa. Raw images of western blots are provided in Supplemental Information Source Data Figure 1a. **(b)** H&E (left) and ER immunohistochemistry (IHC, middle and right inset) are shown for the original patient tumor (top) and HCI-EC-23 PDX model (bottom) from which the HCI-EC-23 cell line was derived. H&E of the patient tumor demonstrates grade 2 endometrioid endometrial carcinoma while the PDX retains differentiated features. IHC for ER demonstrates that both the patient tumor and PDX are positive for ER expression. Scale bars equal 100 μm, inset scale bars equal 50 μm. **(c)** Morphology of the HCI-EC-23 (left) and Ishikawa (right) cell lines under phase-contrast microscopy shows that HCI-EC-23 grows as an adherent monolayer like Ishikawa, but with additional cytoplasmic projections and cell-cell contacts. Scale bar equals 400 μm. **(d)** Immunofluorescence is shown for the epithelial cell marker EPCAM and nuclear stain DAPI for HCI-EC-23 (left) and Ishikawa (right). Individual channels of EPCAM (top left) and DAPI (bottom left) are shown in greyscale. Individual channels are merged and pseudocolored (EPCAM - magenta, DAPI - cyan, right). Scale bars equal 50 μm. **(e)** Bar plots of estrogen response element (ERE) assays after treatment with 10 nM E2 or 1uM of the active metabolite of tamoxifen, 4OHT, are shown. Points represent individual replicates. Error bars represent standard deviation. **(f)** Bar plots of ERE assays are shown for various passages of HCI-EC-23 treated with 10 nM E2. Significance is to respective DMSO treatment. Points represent individual replicates. Error bars represent standard deviation. * p<0.05, **p<0.01, ***p<0.001, and ****p<0.0001.

In order to generate a new EEC cell line that retains ER expression and hormonal responsiveness, we leveraged a set of PDX models[22]. PDXs were dissociated and then grown on collagen-coated plates in a complex media (described in Methods). A major concern when initially growing these dissociated xenografts is the presence and overgrowth of murine stromal cells that out-compete the human tumor cells. To assess the level of murine cell contamination in these initial outgrowths, we used species-specific PCR amplicon length (ssPAL)[34,35], to determine the fraction of cells that are either human or mouse. ssPAL analysis revealed that two of our dissociated xenografts were almost exclusively murine stromal cells; EC-PDX-32 was >99% murine and EC-PDX-33 was 100% murine (no human DNA detected) **(Figure S1a)**. These two xenografts had an overwhelmingly fibroblast-like appearance and eventually stopped growing in culture with no outgrowth of human clones. In contrast, the EC-PDX-23 dissociation led to an outgrowth that was predominantly human (>99%) as detected by ssPAL **(Figure S1a)**. The resulting line, which we have named HCI-EC-23, was subsequently adapted to standard tissue culture-treated dishes and standard RPMI-1640 media supplemented with 10% FBS. Additional testing by IDEXX BioAnalytics (Columbia, MO, USA) detected no murine content and found the line to be mycoplasma negative **(Figure S1a)**. Short tandem repeat (STR) cell line authentication was performed and yielded a novel signature indicating that HCI-EC-23 is not contaminated with cell lines known in either the IDEXX or DSMZ cell line databases (data not shown).

### HCI-EC-23 is an endometrial cancer cell line that retains estrogen signaling

The primary tumor leading to HCI-EC-23 was obtained from a 66-year-old woman who was diagnosed with a grade 2 stage 1A endometrioid endometrial carcinoma (**Table 1**). Additionally, this primary tumor was microsatellite instability-high (MSI-high) and was positive for expression of ER **(Table 1, Figure 1b)**. The PDX model from which HCI-EC-23 was cloned (EC-PDX-23), retained grade 2 histology, MSI-high, and ER positivity (**Table 1, Figure 1b**). MSI testing of the HCI-EC-23 cell line confirmed an MSI-high status (**Table 1**). The HCI-EC-23 cell line grows as a monolayer exhibiting characteristic epithelial cobblestone-like morphology, similar to the Ishikawa cell line, although the HCI-EC-23 cell line appears to exhibit an increased degree of cytoplasmic projections and cell-cell contacts **(Figure 1c)**. Epithelial cell adhesion molecule (EPCAM) positivity, as assessed by immunofluorescent staining, confirms that HCI-EC-23 is an epithelial cell line similar to Ishikawa **(Figure 1d)**.

**Table 1.** Characteristics of HCI-EC-23 primary tumor and derived models.

| Sample | Age | Grade | Stage | MSI Status | MSI Detail | Estrogen Receptor Status |
| --- | --- | --- | --- | --- | --- | --- |
| Primary Tumor | 66 | 2 | 1A | dMMR | Abnormal MLH1/PMS2 | ERα Positive IHC |
| EC-PDX-23 | N/A | 2 | N/A | MSI-H | Unstable at BAT-26, NR-24, MONO-27/NR-27 | ERα Positive IHC |
| HCI-EC-23 Cell Line | N/A | N/A | N/A | MSI-H | Unstable at BAT-26, BAT-25, NR-24, MONO-27/NR-27 | ERα Positive western blot |

MSI = microsatellite instability, dMMR = defective mismatch repair, MSI-H = microsatellite instable high, IHC = immunohistochemistry, N/A = not applicable.

Unlike many of the EEC cell lines described earlier, HCI-EC-23 retains ER expression as detected by western blot **(Figure 1a)** and qRT-PCR **(Supplemental Figure S1c)**. ER is expressed in HCI-EC-23 at approximately half the amount detected in Ishikawa at both the RNA and protein levels **(Supplemental Figure S1c and Figure 1a, respectively)**. ER is also upregulated at the RNA level in HCI-EC-23 upon E2 treatment, while it is repressed in Ishikawa in response to E2 **(Supplemental Figure S1c)**.

We next determined if HCI-EC-23 was responsive to estrogen, similar to the known hormonally responsive cell line Ishikawa. Using a luciferase reporter under the control of tandem estrogen response elements (ERE), ER’s preferred binding site, we found that HCI-EC-23 is indeed responsive to treatment with 10 nM E2 in a manner and magnitude similar to Ishikawa **(Figure 1e**). HCI-EC-23 retains its E2 responsiveness at least through passage 35 **(Figure 1f)**, suggesting it durably remains hormonally responsive in culture.

It is known that the selective ER modulator tamoxifen, which is used to block ER activity for breast cancer treatment, is paradoxically an ER agonist in the normal endometrium and endometrial cancer cells, including the Ishikawa cell line[16, 17, 36, 37]. We found that treatment with 1 μM of the active metabolite of tamoxifen, 4-hydroxy-tamoxifen (4OHT), induces ER activity in both HCI-EC-23 and Ishikawa, demonstrating that these cell lines retain the paradoxical activity of tamoxifen observed in the endometrium **(Figure 1e)**.

## Mutation spectra of HCI-EC-23 is consistent with MSI-high endometrioid endometrial carcinoma

To define the mutational background of the HCI-EC-23 cell line, whole-exome sequencing was performed. We employed a stringent filtering strategy in order to call variants in this cell line as we had no normal sample to remove germline variation; similar strategies have been utilized by the catalog of somatic mutations in cancer (COSMIC) to assess mutations in cell lines[38]. This filtering strategy reduced the number of variants from 29,958 to 3,445, which are presumed to be somatic (**Supplemental File S1**). The filtered, non-silent mutations were largely missense mutations (n = 1,651) followed by frameshift mutations (n = 435) **(Supplemental Figure S2a)**. The mutational profile was largely C >T and most closely matched the mutation signature SBS6, which is associated with defective DNA mismatch repair (cosine-similarity 0.873, **Supplemental Figure S2b**). Next, we compared the mutated genes in HCI-EC-23 to the top twenty most frequently mutated genes in the TCGA EEC UCEC cohort. We found that HCI-EC-23 contained mutations in 11 of the 20 most frequently mutated genes **(Figure 2a)**. Importantly, HCI-EC-23 is mutated at *PTEN, PIK3CA, ARID1A, CTCF*, and *ATM*, but remains wild-type for *PIK3R1, CTNNB1, KRAS*, and *TP53*. Consistent with our goal of establishing a hormonally responsive EEC cell line, the genes that encode progesterone receptor (PR, *PGR*) and ER (*ESR1*) are also wild-type in HCI-EC-23. Copy number alteration (CNA) inference using whole-exome sequencing depth and variant allele fraction (VAF) showed that HCI-EC-23 is relatively copy number neutral aside from the relatively common[5] amplification of chromosome 1q and a few focal alterations (**Figure 2b)**. The mutational landscape, CNA profile, absence of *POLE* exonuclease domain mutation, and MSI-high status demonstrate that HCI-EC-23 is an MSI hypermutated cell line according to the molecular classification identified in the TCGA analysis[5] of endometrial cancer.

**Figure 2.**
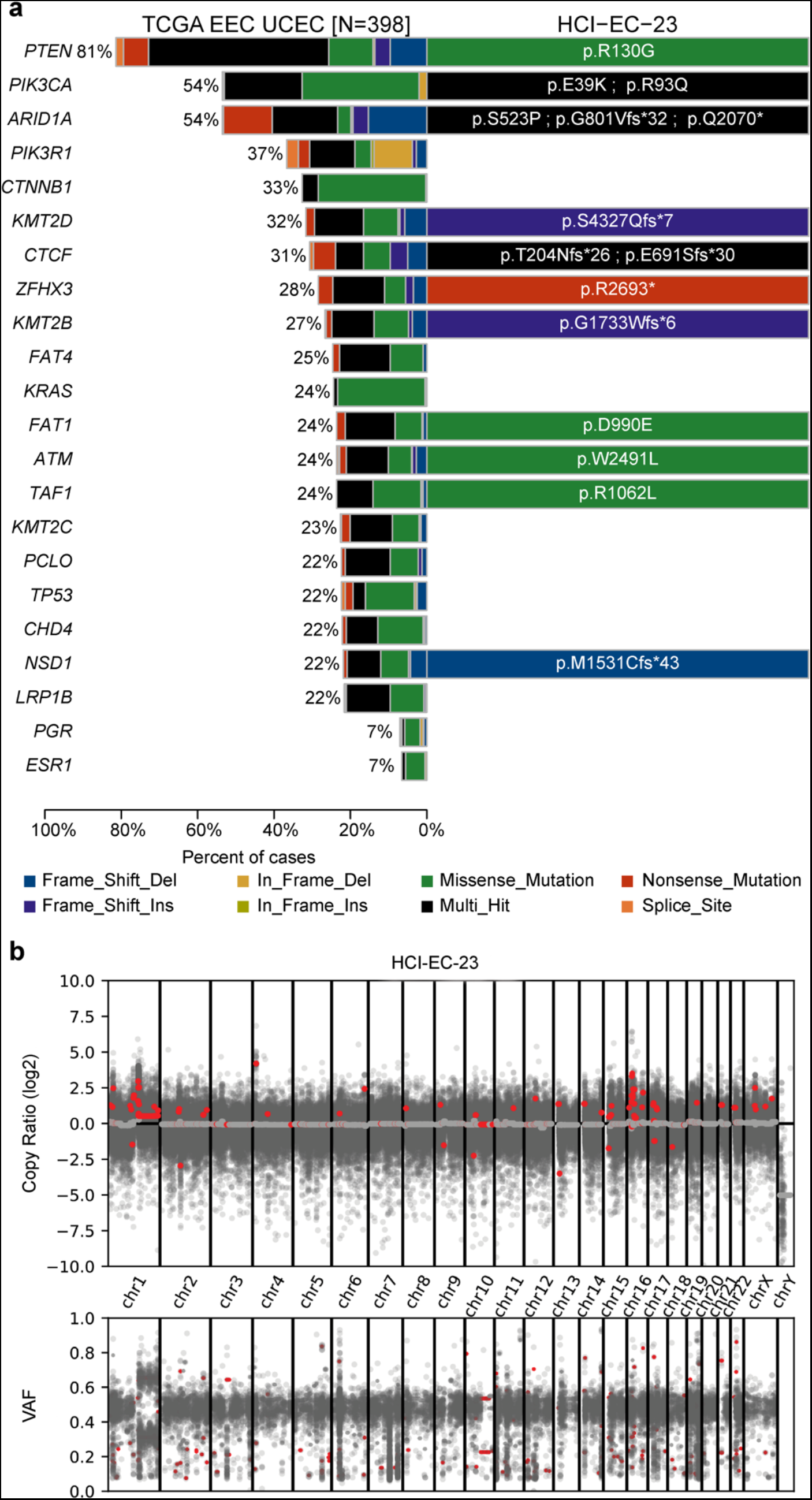
Whole-exome sequencing of HCI-EC-23. **(a)** Displayed is a comparison of the 20 most frequently mutated genes in the TCGA EEC UCEC cohort along with the hormone receptors *PGR* and *ESR1* (left) and mutations identified in HCI-EC-23 (right). Shown are the protein alterations for each gene in HCI-EC-23. **(b)** Plot of log2 copy ratio (top) and variant allele frequency (VAF) along the genome for HCI-EC-23 is shown as computed using CNVKit. The median log2 copy ratio for each computed segment is shown in red.

## HCI-EC-23 exhibits hormone-responsive growth characteristics

Knowing that HCI-EC-23 expresses ER, we next examined the *in vitro* growth characteristics of HCI-EC-23 in the presence of E2 alone and in combination with the growth-suppressive hormone P4. Neither HCI-EC-23 or Ishikawa showed increased growth rates upon treatment with 10 nM E2 **(Figure 3a)**. The absence of a statistically significant E2-driven effect on *in vitro* growth rate for Ishikawa has been previously reported and may be related to differing effects between *in vitro* and *in vivo* conditions[21, 22]. Upon concurrent E2 and P4 treatment (10 nM and 100 nM, respectively), we observed growth reduction in HCI-EC-23, but not in Ishikawa cells **(Figure 3b)**. The P4-mediated growth reduction for HCI-EC-23 is consistent with the growth-suppressive effect of progesterone in the endometrium. The lack of inhibitory effect in Ishikawa may be related to five-fold lower levels of PGR expression compared to HCI-EC-23 **(Figure 4e)**.

**Figure 3.**
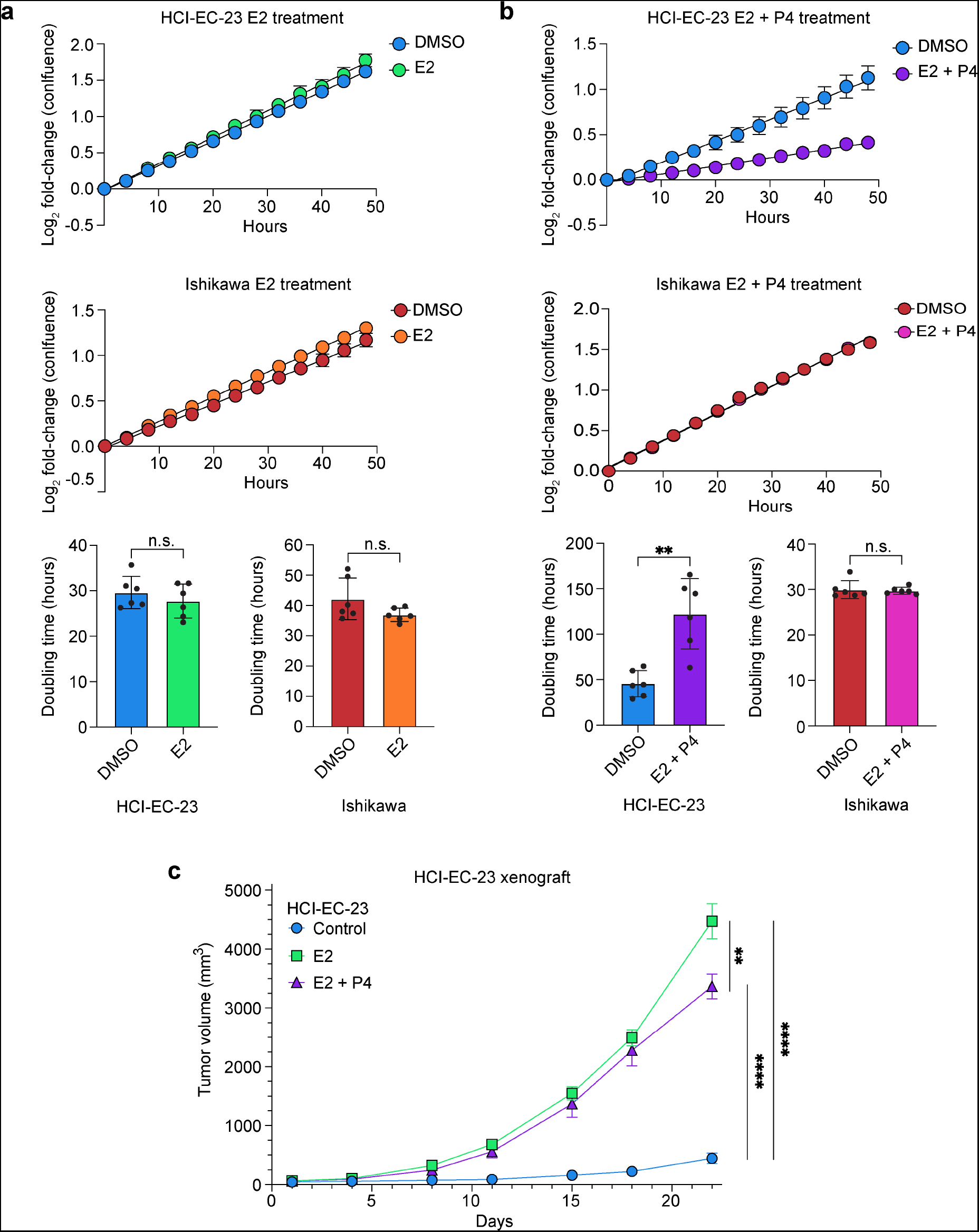
Growth characteristics of HCI-EC-23 *in vitro* and *in vivo*. **(a)***In vitro* log2 growth curves of HCI-EC-23 (top) and Ishikawa (middle) treated with DMSO or 10 nM E2 demonstrate no change in growth rate upon E2 stimulation in both cell lines. Bar plots of doubling times were calculated from these curves and shown for HCI-EC-23 (bottom left) and Ishikawa (bottom right). Data are shown as the mean of six replicates of each cell line. Line graph error bars represent standard error of the mean and barplot error bars represent standard deviation. **(b)***In vitro* log2 growth curves are shown for HCI-EC-23 (top) and Ishikawa (middle) treated with DMSO or 10 nM E2 + 100 nM P4. The addition of P4 results in significant arrest of proliferation in HCI-EC-23. Bar plots of doubling times were calculated from these curves and shown for HCI-EC-23 (bottom left) and Ishikawa (bottom right). Line graph error bars represent standard error of the mean and barplot error bars represent standard deviation. **(c)** Line graph of *in vivo* tumor growth is shown for HCI-EC-23 xenografts in ovariectomized, immunocompromised mice. 1×10^6 cells were injected into the flank of mice treated with E2 (n=4), E2+P4 (n=5), or control (n=5) pellets. Day 1 represents the first day of measurement and was 10 days post-injection. Error bars represent standard error of the mean. *p<0.05, **p<0.01, ***p<0.001, ****p<0.0001, n.s. not significant.

**Figure 4.**
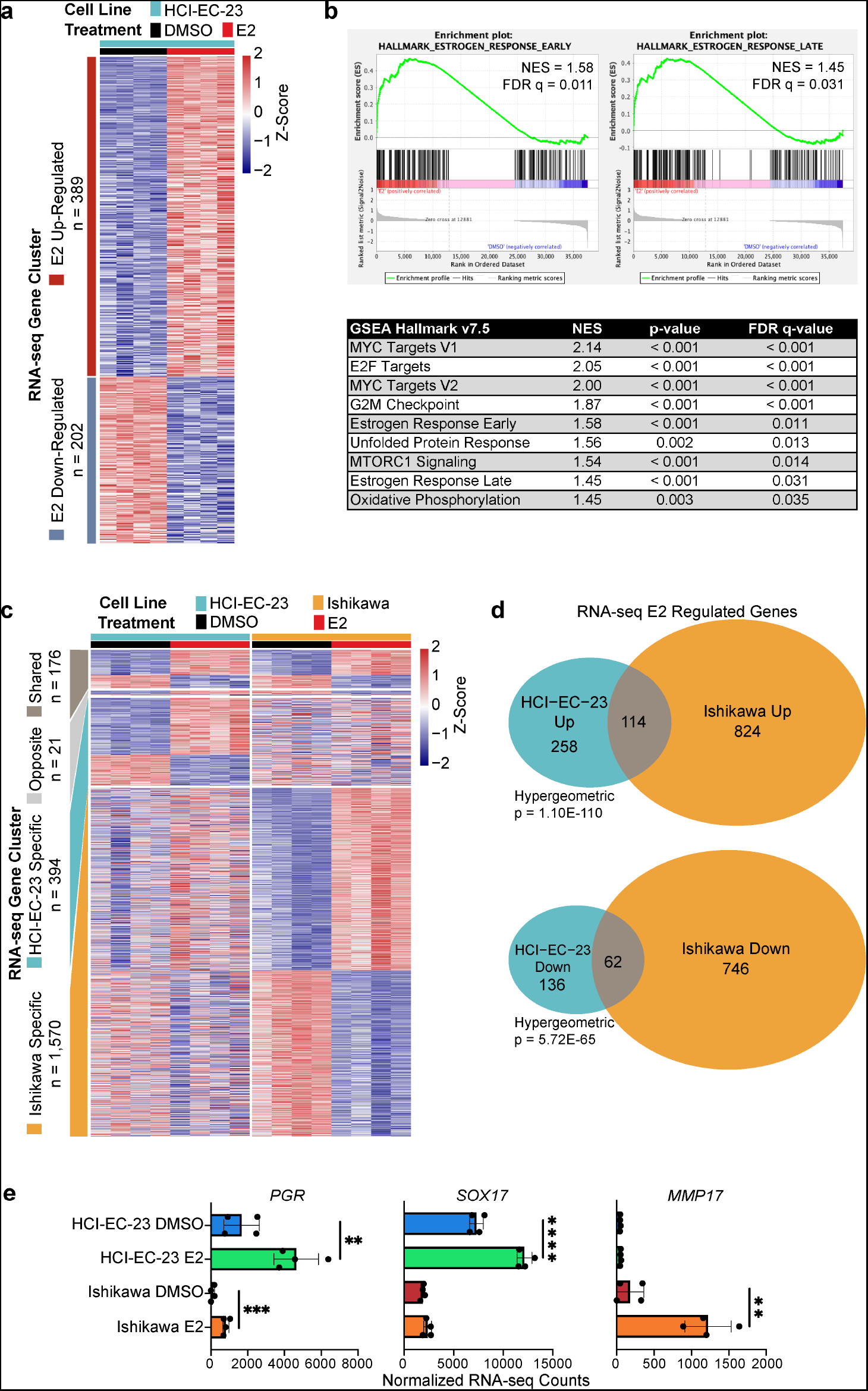
HCI-EC-23 has a transcriptomic response to estrogens that is similar but distinct to Ishikawa. **(a)** A heatmap showing relative expression of genes up- and down-regulated in response to E2 treatment is shown for HCI-EC-23. Each row represents a gene and is annotated by the gene clusters to the left. Samples and treatments are shown by column as indicated in the legend above the heatmap. **(b)** Gene set enrichment analysis (GSEA) plots are shown for the early (left) and late (right) hallmark estrogen response genes. A table of all other significantly enriched hallmark GSEA sets is shown below. NES is the normalized enrichment score. FDR q-value is the false discovery rate. **(c)** A heatmap shows the relative expression of genes up- and down-regulated in response to E2 treatment in HCI-EC-23 compared to Ishikawa cells. Each row represents a gene and is annotated by the gene clusters to the left. Gene clusters identified are shared genes, genes regulated in opposite directions between the two lines, HCI-EC-23 specific, and Ishikawa specific. Samples and treatments are shown by column as indicated in the legend above the heatmap. **(d)** A Venn diagram showing the overlap between E2 up-(top) and down-(bottom) regulated genes in HCI-EC-23 and Ishikawa cell lines. Shown is the hypergeometric p-value for significance of overlap. **(e)** Bar plots are shown for example E2 regulated genes in HCI-EC-23 and Ishikawa. Points represent individual replicates. Error bars represent standard deviation.

We next assessed the growth of HCI-EC-23 *in vivo*. HCI-EC-23 cells were implanted subcutaneously in immunocompromised mice that were ovariectomized to prevent endogenous estrogen production. Mice were also implanted with pellets containing E2, E2 + P4, or vehicle. Individual tumors grew similarly within treatment groups, suggesting that HCI-EC-23 can robustly grow as a xenografted cell line **(Supplemental Figure S3)**. HCI-EC-23 tumors in mice treated with E2 grew substantially faster than tumors in vehicle treated mice; tumors in vehicle treated mice exhibited a very slow growth rate that was 10-fold lower than tumors in E2 treated mice **(Figure 3c)**. These results demonstrate a strong estrogen-dependence for growth *in vivo*, similar growth characteristics have been observed for Ishikawa[22] which far outpaces the effects observed *in vitro*. Tumors in mice treated with E2 + P4 grew similarly to tumors in E2 treated mice at first, but tumor volume began to diverge towards the end of the experiment, where E2 treated tumors were significantly larger than E2 + P4 treated tumors **(Figure 3c)**. This suggests that progesterone impedes *in vivo* growth at later time points, similar to its impact on *in vitro* growth. Combined, the *in vitro* and *in vivo* growth characteristics of HCI-EC-23 demonstrate increased growth upon E2 treatment, especially *in vivo*, while P4 reduced E2 driven growth, consistent with the action of these hormones in the endometrium.

## HCI-EC-23 exhibits a global estrogen response similar to Ishikawa

To assess the transcriptional response to estrogen in HCI-EC-23, we performed RNA-seq following E2 induction. We identified 591 differentially expressed genes following E2 treatment, 389 upregulated and 202 downregulated **(Figure 4a)**. Gene set enrichment analysis (GSEA) demonstrated that both early (NES = 1.58, FDR q-value = 0.011) and late (NES = 1.45, FDR q-value = 0.031) hallmark estrogen responses are upregulated in HCI-EC-23 cells treated with E2, further supporting the estrogen-responsive nature of the cell line **(Figure 4b)**. GSEA also revealed upregulation of MYC targets, E2F targets, G2M checkpoint, unfolded protein response, MTORC1 signaling, and oxidative phosphorylation gene sets **(Figure 4b, Supplemental Figure S4a)**.

We compared the E2 response in HCI-EC-23 and Ishikawa cells. Analysis of RNA-seq data identified four distinct classes of differentially expressed genes in the two cell lines upon E2 induction: 1) shared E2-responsive genes (n = 176), 2) genes regulated in opposite directions (n = 21), 3) genes that are specific to HCI-EC-23 (n = 394), and 4) genes that are specific to Ishikawa (n = 1570) **(Figure 4c, Supplemental Table S1)**. Although the Ishikawa cell line had far more E2-regulated genes than the HCI-EC-23 cell line, we observed statistically significant overlaps between genes upregulated (n = 114, hypergeometric p-value = 1.10E-110) and downregulated (n = 62, hypergeometric p-value = 5.72E-65) in both cell lines, suggesting that they share a core estrogen response **(Figure 4d)**. *PGR*, a known ER target gene, was significantly upregulated in both cell lines **(Figure 4e)**. The two cell lines exhibited a majority of genes whose differential expression was cell line specific. For example, *SOX17*, a transcription factor essential for endometrial gland formation[39–41], was upregulated upon E2 induction in HCI-EC-23, while Ishikawa exhibited lower *SOX17* expression that was not affected by E2 treatment **(Figure 4e)**. Conversely, *MMP17*, a matrix metallopeptidase implicated in extracellular matrix degradation, was upregulated in the Ishikawa cell line, consistent with previous work[21, 42], but was very lowly expressed in HCI-EC-23 **(Figure 4e)**. Together, these data suggest that HCI-EC-23 and Ishikawa share a core estrogen response along with a larger set of cell line specific estrogen-regulated genes.

We also broadly compared the gene expression profile of the E2 treated HCI-EC-23 and Ishikawa cell lines to that of patient samples from TCGA Uterine Corpus Endometrial Carcinoma (TCGA-UCEC) cohort[5]. By principal component analysis (PCA), we found that the TCGA-UCEC cohort can be broadly clustered by histology with endometrioid and serous tumors clustering separately **(Supplemental Figure S4b)**. The HCI-EC-23 cell line clusters with the endometrioid tumors, while the Ishikawa line clusters near the boundary between endometrioid and serous tumors, suggesting that HCI-EC-23 may retain a more differentiated phenotype than Ishikawa **(Supplemental Figure S4b)**. These data are consistent with higher expression of *PGR* and *SOX17* being associated with more well-differentiated tumors[43–47].

Knowing that HCI-EC-23 has a transcriptional response to estrogen similar to that observed in Ishikawa, we next asked if genomic binding of ER is similar upon E2 induction in the two lines. To assess the genomic binding of ER in these lines, we performed ChIP-seq after E2 induction. We found that ER was bound at 1,542 sites in the HCI-EC-23 cell line and 12,361 sites in the Ishikawa cell line **(Figure 5a)**. We hypothesize that the larger number of E2 regulated genes in Ishikawa compared to HCI-EC-23 **(Figure 4c)** is due to this increased genomic binding of ER in Ishikawa compared to HCI-EC-23. The majority (n = 896, 58.1%) of HCI-EC-23 ER ChIP-seq peaks were shared with Ishikawa **(Figure 5a-b)**. Similar overlap frequencies were found when HCI-EC-23 ER peaks were compared to previously published Ishikawa ER peak sets GSE32465[48] (57.8%) and GSE129805[22] (46.0%). ER binding within the 3’ UTR of *PGR* was unchanged between HCI-EC-23 and Ishikawa, consistent with *PGR* being a shared E2-regulated gene **(Figure 5c)**. In contrast, *SOX17* (E2-regulated in HCI-EC-23) and *MMP17* (E2-regulated in Ishikawa) had unique ER binding at their respective loci that are consistent with their cell line specific expression patterns **(Figure 5c)**. The region of the *MMP17* locus that was uniquely bound by ER in Ishikawa cells has been previously shown to be directly involved in the E2-regulation of *MMP17* expression[42, 49]. To assess the importance of these ER bound sites on E2-regulated gene expression, we analyzed the proximity of E2-regulated genes to ER bound sites in both cell lines and found that E2-regulated genes are significantly closer (Wilcoxon signed rank test, p < 2.2E-16) than genes not regulated by E2 to ER bound sites in either HCI-EC-23 or Ishikawa **(Supplemental Figure S5a)**. The close proximity of E2-regulated genes to ER bound sites compared to genes not regulated by E2 suggests functional importance for many ER bound sites in regulating E2-responsive genes.

**Figure 5.**
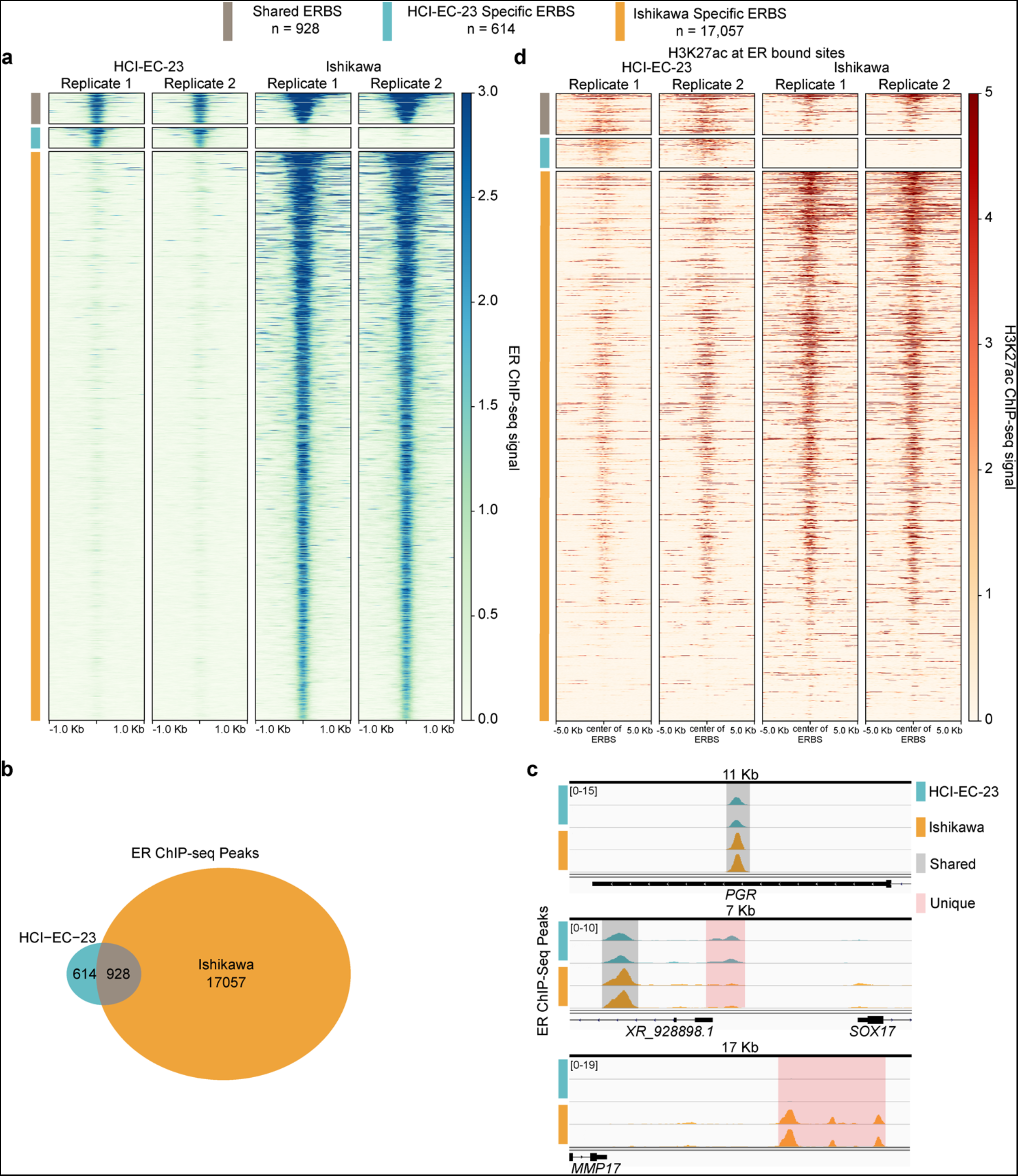
ER genomic binding in HCI-EC-23 is similar but distinct to Ishikawa. **(a)** A heatmap is shown for ER genomic binding signal. Each row represents a ChIP-seq peak and is annotated by estrogen receptor bound site (ERBS) category – either shared ERBS, HCI-EC-23 specific ERBS, or Ishikawa specific ERBS according to the legend above the heatmap. Each column is a sample (two replicates per cell line). **(b)** A Venn diagram shows the overlap between ER ChIP-seq peaks in HCI-EC-23 and Ishikawa cell lines. **(c)** Genome browser tracks plotting ER ChIP-seq peaks are shown for HCI-EC-23 (top) and Ishikawa (bottom). Regions shown are the 3’ UTR of *PGR*, the region upstream of *SOX17*, and the region downstream of *MMP17*. Shared peaks between the two cell lines are highlighted in gray, while unique peaks are highlighted in red. Peak height is scaled within each locus. **(d)** A heatmap shows the H3K27ac signal without E2 treatment at ER bound sites. Each row represents H3K27ac signal centered around an identified ERBS (and is ordered as in panel a). The different categories of ERBS are annotated (as in panel a) according to the legend above the heatmap. Each column is a sample (two replicates per cell line).

As expected, motif analysis identified ESR1 (ER) as the dominant motif in shared, HCI-EC-23-specific, and Ishikawa-specific ChIP-seq peaks **(Supplemental Figure S5b)**. Among shared ER ChIP-seq sites, PITX2 and ZN770/MAF/ZSC22/SMAD3 motifs are also significantly enriched **(Supplemental Figure S5b)**. HCI-EC-23 specific ER ChIP-seq sites were enriched for ATF3 and ZN770/ZSC22/ZFX/CRX motifs, while Ishikawa specific ER ChIP-seq sites were enriched for AP-1 family members and PITX2/ZN264 motifs **(Supplemental Figure S5b)**. These differences in motif enrichment suggest that different transcription factors could contribute to differential ER genomic binding or subsequent effects on transcription.

Looking more closely at ERE sequences within the ER ChIP-seq sites, we found that 61.7% of shared sites contained an ERE sequence as compared to 43.0% of HCI-EC-23 specific sites or 25.6% of Ishikawa specific sites **(Supplemental Figure S5c)**. The frequency of EREs in these binding sites suggests that ER binding in HCI-EC-23 is strongest at sites with frequent EREs as compared to Ishikawa, where more sites are bound that do not harbor EREs. In addition to differences in ERE frequency, differences in chromatin states between HCI-EC-23 and Ishikawa may explain cell line specific ER binding. To interrogate chromatin states at ER bound sites, we assessed H3K27ac ChIP-seq signal in DMSO treated HCI-EC-23 and Ishikawa cells as a baseline marker of active regulatory regions. We found that H3K27ac signal was relatively high in both cell lines at shared ER bound sites **(Figure 5d, Supplemental Figure S5d)**. Conversely, cell line specific ER bound sites are associated with reduced H3K27ac signal in the other line, suggesting that baseline chromatin state is important for cell line specific ER binding **(Figure 5d, Supplemental Figure S5d)**. We also observed differences in the distribution of ER bound sites relative to transcription start sites between HCI-EC-23 and Ishikawa cell lines **(Supplemental Figure S5e)**. ER is bound more at distal intergenic regions in HCI-EC-23 while in Ishikawa ER binding is expanded at promoters and 5’UTRs. Overall, these gene regulation studies indicate that HCI-EC-23 cells exhibit a robust transcriptional response to estrogen that is similar to Ishikawa.

## Discussion

Although most endometrial cancers are hormonally driven and estrogen-responsive, only one currently available endometrial cancer cell line has retained this estrogen-responsiveness, the Ishikawa cell line. The lack of additional cell line models to examine the estrogen response in endometrial cancer has limited our understanding of the mechanisms of pathogenesis and the study of therapeutic interventions. In this study, we described a novel estrogen-responsive cell line, HCI-EC-23, that will benefit the field by providing an additional cell line model to study the hormonally driven aspects of endometrial cancer. Importantly, this model has characteristics that are distinct from the Ishikawa cell line, including sensitivity to progesterone treatment. The sensitivity of HCI-EC-23 to progesterone will be particularly important for translational studies, since synthetic progesterone (progestins) are the only approved pharmaceutical treatment for primary endometrial cancer. We provided our strategy to develop cell lines from PDX tumors along with methods to rapidly determine murine contamination levels by using ssPAL. This strategy led to the isolation of HCI-EC-23, from a PDX model derived from a grade 2, MSI-high, and ER-positive endometrioid endometrial tumor. Critically, HCI-EC-23 retained ER expression as shown by western blot and qRT-PCR and ER was functional as assayed by ERE-driven luciferase. HCI-EC-23 also displays paradoxical activation of ER by tamoxifen, an effect known in the endometrium and Ishikawa cell line[16, 17, 36, 37]. HCI-EC-23 showed a strong P4-induced proliferation arrest *in vitro*. HCI-EC-23 xenografts showed strong E2-dependence *in vivo* with a reduction in tumor volume with co-administration of P4. These results demonstrate that HCI-EC-23 is a translationally relevant estrogen-responsive endometrial cancer cell line.

Whole-exome sequencing and MSI typing provided evidence that HCI-EC-23 harbors mutations to many of the genes frequently mutated in endometrioid endometrial cancer. We also showed that HCI-EC-23 has relatively few copy-number alterations and is MSI-high, placing it into the MSI-high molecular subgroup identified by TCGA. Ishikawa is also MSI-high but harbors mutations distinct from HCI-EC-23. Among the top 20 mutated genes in endometrioid endometrial carcinoma, Ishikawa is mutant for *PIK3R1*, *FAT4*, *KMT2C*, *PCLO*, *TP53*, and *LRP1B*[50]. In contrast, HCI-EC-23 is mutant for *PIK3CA*, *KMT2B*, *TAF1*, and *NSD1*. Both lines are mutant for *PTEN*, *ARID1A*, *KMT2D*, *CTCF*, *ZFHX3*, *FAT1*, and *ATM*, although the specific alterations to these genes are different between the two lines[50]. This demonstrates that HCI-EC-23 and Ishikawa have similar but distinct genetic backgrounds. For instance, both cell lines harbor two common alterations to the PI3K pathway, sharing altered *PTEN* but differ in having either mutant *PIK3CA* or *PIK3R1*. Used together, the similar but distinct genetic backgrounds between the two lines are more reflective of the mutational heterogeneity of patient tumors when studying the hormonal aspects of endometrial cancer than either line alone.

Molecular characterization of the estrogen response using RNA-seq and ER ChIP-seq in HCI-EC-23 showed many parallels to Ishikawa. A large core set of genes were up- or down-regulated upon E2 induction in both HCI-EC-23 and Ishikawa. This coordinated regulation in both cell lines may be attributable to the significant overlap of ER binding sites in the two lines. However, there remain estrogen-regulated genes that are unique to each cell line. This difference may be due to the different genetic backgrounds and chromatin states of the two lines. Alternatively, the relatively larger number of ER binding sites in Ishikawa may confer regulation of a proportionally larger set of genes, which is likely related to the higher expression of ER in Ishikawa compared to HCI-EC-23. Differences in the degree of cellular differentiation in the two lines could also alter the set of estrogen-regulated genes, as suggested by RNA-seq clustering that indicates HCI-EC-23 is more similar to endometrioid TCGA tumors while Ishikawa clusters among a subset of endometrioid tumors closer to serous tumors. HCI-EC-23 expresses higher levels of both progesterone receptor and SOX17 than Ishikawa, both of which are transcription factors essential for endometrial gland development and function[39–41, 51, 52]. In the context of endometrial cancer, both progesterone receptor and SOX17 are tumor suppressors and exhibit reduced expression that is associated with less differentiated, high-grade tumors[43–47]. Progesterone receptor and SOX17 have also been shown to cooperatively regulate target genes as part of a transcriptional network[53–55]. Together, these data suggest that HCI-EC-23 cells may have a more differentiated state than Ishikawa.

As HCI-EC-23 is an adherent cell line, it is more easily amenable to screening efforts and genetic manipulation than PDX models due to the large numbers of cells and conditions required. Although the power of PDX models to recapitulate tumor heterogeneity is clear, the hormonal aspects of these models are often less clear and are largely only assessed by hormone-receptor positivity[30–33]. We believe that HCI-EC-23 is uniquely situated to vastly improve the quality of research regarding the molecular aspects of hormonally driven endometrial cancer by providing another well-characterized cell line model for investigators to utilize.

## Methods

### Endometrial cancer patient-derived xenograft (PDX) models

PDX models were generated from human tissue acquired with patient informed consent and experiments performed in accordance with University of Utah Institutional Review Board approved guidelines and regulations[22]. All animal procedures were conducted following approved University of Utah Institutional Animal Care and Use Committee guidelines and regulations. Animals were housed in the Comparative Medicine Center at the University of Utah under standard conditions. Tumor specimens from patients with endometrial cancer were collected in the operating room by the Biorepository and Molecular Pathology Shared Resource. Tumor specimens were transported in ice-cold RPMI-1640 base medium to the lab. Each tumor specimen was mechanically divided into multiple pieces using sterile disposable scalpels. A minimum of three female 6- to 8- week old nude mice (nu/nu, Jackson Laboratory) were used for tumor implantation. Orthotopic implantation of ~4 mm^3^ tumor fragments into the lumen of the mouse uterus was performed as described previously[22]. Nude mice were not given supplemental estrogen. Tumor size was measured twice per week. Mice were sacrificed and necropsies performed once tumor size reached 2 cm in diameter.

### Immunohistochemical staining of tumor specimens

Immunohistochemical staining of FFPE slides was performed by ARUP Laboratories using an automated immunostainer. Briefly, 5 μm tissue sections were obtained using a standard microtome and de-paraffinized with the EZ Prep solution. The slides were then treated with Cell Conditioning 1 (CC1), pH 8.5 for 68 min at 95°C. The primary antibody (#ab80922, Abcam) was applied for 1 hr at a dilution of 1:100 at 35°C. Following removal of the primary antibody, the secondary antibody (#B8895, Sigma-Aldrich) was applied for 1 hr at a dilution of 1:100 at 37°C. Slides were then exposed to the IView DAB Map detection kit (Ventana) and counterstained with hematoxylin for 8 min. After dehydration in graded alcohol, coverslips were put on and slides were read by board-certified pathologists. Hematoxylin and eosin (H&E) staining was performed on 5 μm tissue sections.

### Tumor dissociation

Patient-derived xenograft (PDX) samples were cut into approximately 2 – 4mm chunks with forceps and a scalpel and transferred to a gentleMACS C tube (Miltenyi Biotec, 130-093-237) containing 4.7 ml RPMI 1640 base media, 200 ml of Enzyme H, 100 ml of Enzyme R and 25 ml of Enzyme A (Miltenyi Biotec, Tumor Dissociation Kit 130-095-929). Tumor samples were dissociated using the gentleMACS h_tumor_01 program for 1 hour at room temperature. The sample was resuspended and filtered through a 70 mm MACS SmartStrainer (Miltenyi Biotec, 130-098-462) and rinsed with 20 mL of RPMI 1640 media (Gibco, 11875135). The single-cell suspension was centrifuged at 300g for 7 minutes. The supernatant was aspirated and cells were resuspended, counted, plated in patient-derived xenograft organoid (PDXO) media, and grown at 37°C in 5% CO_2_. PDXO media consists of Advanced DMEM/F12 (Gibco, 12634010) supplemented with 5% heat-inactivated FBS (Thermo Fisher), 10% Pen-Strep (Gibco, 15140148), 1x HEPES (Gibco, 15630080), GlutaMAX Supplement (Gibco, 35050061), hEGF (Sigma Aldrich, E9644), 10 mM Y-27632 (Selleckchem, S1049), 10 ng/ml FGF-b/FGF-2 (Invitrogen, 68-8785-82), and 1 mM N-Acetyl-L-cysteine (Sigma Aldrich, A7250).

### Cell culture

The established endometrial carcinoma cell lines were obtained from ATCC (AN3CA[10], HEC-1-A[11], and RL95-2[12]), Sigma-Aldrich (Ishikawa[8, 9], MFE-280[13], MFE-296[13]), or Riken (JHUAS-1[14] and JHUEM-2[14]). Ishikawa was cultured in RPMI 1640 (Gibco, 11875119) supplemented with 10% fetal bovine serum (Gibco, 26140079) and 1% penicillin-streptomycin (Gibco, 15140163). AN3CA, RL95-2, HEC-1-A, JHUEM-2, and JHUAS-1 were cultured in DMEM/F12 (Gibco, 11320033) supplemented with 10 % fetal bovine serum (Gibco, 26140079) and 1% penicillin-streptomycin (Gibco, 15140163). MFE-296 was cultured in MEM (Gibco, 11095072) supplemented with 10% fetal bovine serum (Gibco, 26140079) and 1% penicillin-streptomycin (Gibco, 15140163). MFE-280 was cultured in MEM (Gibco, 11095072) supplemented with 10% fetal bovine serum (Gibco, 26140079), 1% insulin-transferrin-selenium (Gibco, 41400045), and 1% penicillin-streptomycin (Gibco, 15140163). The HCI-EC-23 cell line was initially grown in PDXO media (described in tumor dissociation section) and were adapted to RPMI 1640 (Gibco, 11875119) supplemented with 10% fetal bovine serum (Gibco, 26140079) and 1% penicillin-streptomycin (Gibco, 15140163). All cells were cultured at 37°C in 5% CO_2_.

For experiments with hormone treatments, cells were cultured in hormone deprived media instead of the regular media (described above). Hormone deprived media consisted of phenol red-free RPMI1640 (Gibco, 11835055) supplemented with 10% charcoal-stripped fetal bovine serum (Sigma Aldrich, F6765) and 1% penicillin-streptomycin (Gibco, 15140163). Hormones used were 17β-estradiol (E2, Sigma Aldrich, E2758), (Z)-4-Hydroxytamoxifen (4OHT, Sigma Aldrich, H7904), and progesterone (P4, Sigma Aldrich, P3972). Hormones were dissolved in dimethyl sulfoxide (DMSO, Fisher Scientific, BP231-100).

### ssPAL (species-specific PCR amplicon length)

ssPAL was performed as described previously[34, 35]. Primer sets used were reported in Schneeberger et al. 2016[35]. Primer set 5 (Forward: TCATTGGCTTAAAATGTGT, Reverse: FAM-TTTATTTTAAGGGGTTGTAATG) amplifies a region of Ribonuclease P, yielding a 272 nt amplicon in humans and a 278 nt amplicon in mice. Primer set 43 (Forward: CTATTCCTATAGCACAAAGG, Reverse: FAM-GATGGTGTACACCCATCATG) amplifies a region downstream of *RC3H2*, yielding a 211 nt amplicon in humans and a 206 nt amplicon in mice. PCR was performed using 2X Phusion HF Mastermix (NEB M0531L) in 10 μL reactions with 20 ng of template DNA. Cycling conditions were 98°C 5 minutes; 30 cycles of 98°C 30 seconds, 52°C 30 seconds, 72°C 30 seconds; 72°C 10 minutes. Samples were diluted 1:20 and then mixed 1:1 by DNA source allowing for analysis of multiple assays for one specimen at a time. Samples were then sent to GeneWiz (South Plainfield, NJ, USA) for fragment analysis by capillary electrophoresis. Fragment traces were then analyzed using the Peak Scanner module on ThermoFisher Cloud. Peaks were called human in the following ranges (primer set 5 271-273 nt; primer set 43 210-212 nt) and mouse (primer set 5 278-279 nt; primer set 43 205-207 nt). Percent human or mouse DNA was calculated by dividing the area of the human or mouse peak by the sum of both peaks.

### MSI testing

Microsatellite Instability (MSI) testing was assessed at the quasimonomorphic mononucleotide runs[56] BAT-26, BAT-25, NR-21, NR-24, and NR-27/MONO-27 by the University of Utah Health Sciences Center Genomics Core. Fragment analysis of fluorescently labeled PCR products was performed on the Applied Biosystems 3730 platform. Primers utilized were BAT-26 (Forward: 6FAM-CTGCGGTAATCAAGTTTTTAG, Reverse: gtttctAACCATTCAACATTTTTAACCC), BAT-25 (Forward: VIC-TACCAGGTGGCAAAGGGCA, Reverse: gtttctTCTGCATTTTAACTATGGCTC), NR-21 (Forward: NED-GAGTCGCTGGCACAGTTCTA, Reverse: gtttctCTGGTCACTCGCGTTTACAA), NR-24 (Forward: PET-GCTGAATTTTACCTCCTGAC, Reverse: gtttctATTGTGCCATTGCATTCCAA), NR-27/MONO-27 (Forward: 6FAM-AACCATGCTTGCAAACCACT, Reverse: gtttctCGATAATACTAGCAATGACC). Sites were considered contracted or unstable if they varied by >5 nt from a reference normal sample.

### ERE (estrogen response element) luciferase reporter assay

HCI-EC-23 and Ishikawa cells were grown in hormone-deprived media for 3 days prior to plating at 15,000 cells per well in a 96-well plate in hormone-deprived media. One day after plating, cells were transfected, according to manufacturer’s protocol, the following day with dual ERE-driven firefly luciferase and constitutive *renilla* luciferase constructs (Qiagen, CCS-005L) using FuGENE HD transfection reagent (Promega, E2312). One day post-transfection, media was changed and cells were treated with either DMSO, 10 nM E2, or indicated concentrations of 4OHT in hormone-deprived media. Twenty-four hours post-treatment, luciferase activity was measured using the Dual-Luciferase Reporter assay system (Promega, E1960) on a Promega Glo-Max luminometer (Promega, 9301-010). All experiments were performed in technical triplicate with three biological replicates. Firefly luciferase was normalized to *renilla* luciferase and then fold change determined to respective DMSO control. Statistical significance was determined by Student’s T-test for comparison of treatments to control DMSO.

### Cell proliferation assays

Ishikawa and HCI-EC-023 cells were initially cultured in regular media and transferred to hormone-deprived media 48 hours prior to plating for proliferation experiments. For proliferation assays, 96-well plates were seeded with 10,000 cells per well and incubated overnight at 37°C. The following day, control DMSO, 10 nM estrogen, or the combination of 10 nM E2 and 100 nM P4 were added to wells, after which the plates were loaded onto the IncuCyte ZOOM live cell imaging system (Sartorius, 4459). Cell proliferation was monitored via time-lapse image acquisition (2 images per well) every 4 hours for up to 96 hours using the IncuCyte(R) ZOOM software (Sartorius, vers. 2016B, 2018A). The initial acquisition was used as 0-hour timepoint. Doubling times were calculated by first normalizing confluency measurements to cell density at 0-hour timepoint. The log_2_ values of each normalized confluency measurement were then calculated, and the first 48 hours of each growth curve were fitted to a linear regression model. Doubling times were determined using 1/slope of each best fit line, and statistical significance was assessed by Student’s T-test using GraphPad Prism. Cell proliferation assays were performed as two biologic replicates assayed in technical triplicate.

### qRT-PCR (quantitative reverse-transcription polymerase chain reaction)

HCI-EC-23 and Ishikawa cells were grown in hormone-deprived media for five days. Total RNA was harvested following an 8 hour DMSO or 10 nM E2 induction with buffer RLT plus (Qiagen, 1053393) supplemented with 1% beta-mercaptoethanol (Sigma Aldrich, M3148-100ML). Total RNA was purified using a Quick-RNA Miniprep kit (Zymo Research, R1055). Twenty-five nanograms of RNA was used as starting material per sample for qRT-PCR using the Power SYBR Green RNA-to-Ct 1-Step kit (Thermo Scientific, 4389986) on a Bio-Rad CFX96 thermocycler. Expression measurements were calculated using the ΔΔCt method using *CTCF* as a control. Primers used were for *ESR1* (Forward: GGGAAGTATGGCTATGGAATCTG, Reverse: TGGCTGGACACATATAGTCGTT) and *CTCF* (Forward: ACCTGTTCCTGTGACTGTACC, Reverse: ATGGGTTCACTTTCCGCAAGG). Experiments were performed in biologic triplicate and statistical significance was determined by Student’s T-test to respective DMSO control.

### Western blotting

HCI-EC-23, Ishikawa, AN3CA, HEC-1-A, JHUEM-2, MFE-280, MFE-296, JHUAS-1, and RL95-2 cells were grown in regular media (described in cell culture methods) to approximately 80% confluence in 10-cm dishes. Cells were washed with PBS, trypsinized, quenched with media, and centrifuged at 200xg for 5 minutes to pellet cells. Cell pellets were washed with ice-cold PBS, centrifuged at 200xg for 5 minutes, PBS aspirated, and kept on ice. Cell pellets were lysed with 3-5x packed cell volume of RIPA buffer (1x PBS, 1% NP-40, 0.5% sodium deoxycholate, 0.1% SDS) supplemented with protease and phosphatase inhibitors (Thermo Scientific, A32959). Cells were lysed on ice for 30 minutes with periodic vortexing every 10 minutes. Lysates were cleared by centrifugation at 12,000xg for 15 minutes at 4°C. The cleared supernatant was kept on ice and protein was quantified using the Qubit Protein Assay kit (Thermo Scientific, Q33212) and measured on the Qubit 2.0 Fluormeter (Thermo Scientific, Q32866). Li-COR chameleon duo pre-stained protein ladder (Li-COR, 928-60000) along with 100 μg of protein combined with loading buffer (Thermo Scientific, NP0007) were loaded onto a 4%-12% Bis-Tris gel (Thermo Scientific, NP0336BOX) and run at 150 V in 1x MOPS SDS running buffer (Thermo Scientific, NP0001). Protein was then transferred from the gel onto a polyvinylidene difluoride (PVDF) membrane (Thermo Scientific, IB24002) using the iBlot 2 dry blotting system (Thermo Scientific, IB21001). The PVDF membrane was blocked in Intercept TBS blocking buffer (Li-COR, 927-60001) for 1 hour at room temperature. Blots were then probed simultaneously with primary antibodies to rabbit anti-ERα (Santa Cruz Biotechnology, sc-543) and mouse anti-β-Actin (Santa Cruz Biotechnology, sc-47778) at 1:1,000 dilution in Intercept TBS blocking buffer with 0.1% Tween-20 overnight at 4°C. Blots were washed 3x for 10 minutes with 1x TBS-T (1x TBS, 0.1% Tween-20) and then probed simultaneously with secondary antibodies IRDye 800CW donkey anti-rabbit (Li-COR, 926-32213) and IRDye 680RD donkey anti-mouse (Li-COR, 926-68072) at 1:10,000 dilution in Intercept TBS blocking buffer with 0.1% Tween-20 for 1 hour at room temperature. Fluorescence signal from the antibodies were then detected using the Azure 600 imaging system (Azure Biosystems, AZI600-01) with the 800CW and 680RD channels. Western blot images were analyzed using Image Studio Lite software (Li-COR). Raw western blot images are provided in Supplemental Figures Source Data Figure 1a.

### Whole-exome sequencing

Whole-exome sequencing was performed on the HCI-EC-23 cell line using the Agilent SureSelect XT Human All Exon v7 capture kit. The library was then sequenced on an Illumina NovaSeq 6000 yielding 150 bp paired-end reads. The Whole Exome Sequencing – BWA + GATK 4.0 pipeline (https://cgc.sbgenomics.com/public/apps/admin/sbg-public-data/whole-exome-sequencing-bwa-gatk-4-0/41) was then utilized on SevenBridges Genomics to align reads to hg38 and generate a variant call file (VCF). The SevenBridges pipeline yielded 162,787,374 mapped reads leading to an overall 96.01% of targeted bases having at least 30X coverage as determined by Picard (https://broadinstitute.github.io/picard/) CollectHsMetrics. VCF2MAF (https://github.com/mskcc/vcf2maf) was used to convert the VCF file to a mutation annotation file (MAF) while also annotating variant effects using the Ensembl Variant Effect Predictor[57] (VEP) v102. This MAF file contained 29,958 variants and was filtered to remove presumed germline mutations using the following strategy. All variants tagged as a “common variant” (defined as an allele frequency in at least one gnomAD[58] subpopulation >0.04%) were removed. The remaining variants were filtered to include those without existing variant identifiers, present in COSMIC[38], while removing those with identifiers in dbSNP[59] with any reported allele frequency. This resulted in a filtered MAF file containing only 3,445 mutations which are presumed to be somatic. The MAFTools[60] R package was used to generate mutational frequency, mutation-type, and mutational signature plots. Mutational signature plots were made using the mutational profiles outlined in Alexandrov et. al. 2020[61]. MAFTools also contained a MAF file of the TCGA uterine corpus endometrial cancer (UCEC) cohort[5, 62] that was filtered for endometrioid histology and used to generate the barplot comparing mutational frequency in TCGA to HCI-EC-23 mutations. CNVkit[63] was used to assess copy number alterations (CNA) from the whole-exome sequencing. Because of the lack of paired-normal DNA, CNA analysis with CNVkit was performed against a flat reference of the hg38 genome. The scatter function in CNVkit was used to generate a plot of computed CNA in the HCI-EC-23 cell line along with a plot of VAF using the VCF from the SevenBridges pipeline (described above).

### RNA-seq

HCI-EC-23 or Ishikawa cell lines were cultured in hormone-deprived media for five days and then induced with 10 nM E2 or DMSO (vehicle) for eight hours. RNA was harvested by washing cells with PBS and lysing cells with buffer RLT plus (Qiagen, 1053393) supplemented with 1% beta-mercaptoethanol (Sigma-Aldrich, M3148). RNA was then purified using the Zymo Quick-RNA Miniprep kit (Zymo Research, R1055). RNA-seq libraries were generated using the KAPA Stranded mRNA HyperPrep kit (Roche, KK8580/08098115702) using 500 ng of RNA. RNA-seq libraries were pooled and sequenced on either the Illumina HiSeq 2500 or NovaSeq 6000. Two replicates of each cell line were sequenced on each sequencing platform. Sequencing reads were aligned to the hg38 build of the human genome using HISAT2[64]. SAMtools[65] was used to convert SAM files to BAM files. Genes were defined by the University of California Santa Cruz (UCSC) Known Genes and reads mapped to known genes were assigned with featureCounts[66]. Within each cell line, read counts were normalized and analyzed for differential expression via the DESeq2[67] package in R[68] while controlling for the sequencing platform by using the design formula (= ~Sequencer + Treatment), where sequencer reflects the sequencing platform and treatment is 10 nM E2 or DMSO. Heatmaps of gene expression z-score by gene were made using the pheatmap R package.

### The Cancer Genome Atlas UCEC RNA-sequencing analysis

The Cancer Genome Atlas (TCGA) RNA-seq dataset for Uterine Corpus Endometrial Carcinoma (TCGA, PanCancer Atlas) was downloaded from Xena browser (https://gdc-hub.s3.us-east-1.amazonaws.com/download/TCGA-UCEC.htseq_counts.tsv.gz). Histological data for the TCGA samples was obtained from cBioPortal[69] (https://www.cbioportal.org/study/summary?id=ucec_tcga_pan_can_atlas_2018). Counts were converted from log(2+1) to raw counts. Counts for E2 treated Ishikawa and HCI-EC-23 samples were determined as described in the RNA-seq section. These raw counts were then combined with the TCGA samples and analyzed using DESeq2[67]. Counts were corrected for their source using the design formula (=~Group). Group=Gertz vs TCGA. Batch effects of the different groups were removed from the variance stabilized counts using the limma[70] package function removeBatchEffect(). Normalized counts were then used to create a PCA plot.

### ChIP-seq

HCI-EC-23 and Ishikawa cells were grown in hormone-deprived media for 5 days prior to collection. For ER ChIPs, cells were induced with 10 nM E2 for 1 hour. For H3K27ac ChIPs, cells were induced with DMSO for 8 hours. Cells were then fixed using 1% formaldehyde (Sigma Aldrich, 252549) for 10 minutes at room temperature to cross-link DNA and proteins. Cross-linking was stopped by addition of glycine (Fisher Scientific, BP381) to a final concentration of 125 mM. Cells were then washed with cold PBS and harvested via scraping in Farnham lysis buffer supplemented with protease and phosphatase inhibitors (Thermo Scientific, A32959). Farnham lysis buffer consisted of 5 mM PIPES pH 8.02, 85 mM KCl, and 0.5% NP-40. Chromatin immunoprecipitation was performed as previously described[71] with 5 μg of an Anti-ERα (Millipore Sigma, 06-935) or Anti-H3K27ac (Active Motif, 39133) antibody. ChIP-seq libraries were sequenced on an Illumina NovaSeq 6000. Sequencing reads were aligned to the hg38 build of the human genome using Bowtie[72] using parameters: -m 1 -t –best -q -S -I 32 -e 80 -n 2. MACS2[73] was used to call peaks with a *P*-value cutoff of 1×10^− 10^ and mfold parameter between 15 and 100. Input control libraries from either cell line were used as controls for each ChIP-seq experiment. All ChIP-seq experiments were performed in biologic duplicates. Overlaps between peaks were determined using a 1-bp minimum overlap using BEDTools[74]. Heatmaps and profile plots of ChIP signal were made using deepTools[75]. Motif discovery was performed on 500-bp regions surrounding the summit of identified peaks. Motifs between 6 and 30 bp in size were identified by MEME-ChIP, part of the MEME suite[76], with a motif distribution of zero to one occurrence per sequence. Percentage of peaks containing EREs was identified using AME, part of the MEME suite[76], with the ESR1_HUMAN.H11MO.0.A position weight matrix from HOCOMOCOv11[77]. Genomic annotation of binding sites was performed using CEAS[78].

Previously published Ishikawa ER ChIP-seq peak calls (narrowPeak or broadPeak files) were downloaded from GEO datasets GSE32465[48] (sample GSM803422) and GSE129805[22] (samples GSM3722307 and GSM3722311). Overlaps between individual replicates were determined as described above and peaks were remapped from hg19 to hg38 using UCSC Genome Browser’s LiftOver tool. These data were then used to identify overlapping Ishikawa ER peaks with HCI-EC-23 ER peaks with BEDTools.

### Immunofluorescence

HCI-EC-23 and Ishikawa cells were grown in regular media in 8-well chamber slides (Thermo Scientific, 177402PK). Cells were washed with 1x PBS and then fixed using 4% formaldehyde (Sigma Aldrich, 252549) in 1x PBS for 10 minutes at room temperature. After fixation, cells were washed 3x with ice-cold 1x PBS-T (1x PBS, 0.1% Tween-20). Cells were then permeabilized with Triton X-100 (Sigma Aldrich, T8787) diluted in 1x PBS (1x PBS, 0.1% Triton X-100) for 10 minutes at room temperature. Cells were then washed 3x for 5 minutes with 1x PBS-T. Cells were then blocked using a BSA blocking buffer (1x PBS-T, 1% BSA, 22.5mg/mL glycine) for 30 minutes at room temperature. Blocking buffer was then removed and cells were incubated with an APC-conjugated anti-human EpCAM (BioLegend, 369809) antibody diluted 1:20 in dilution buffer (1x PBS-T, 1% BSA) for 1 hour at room temperature in the dark. Cells were then washed 3x 5 minutes in 1x PBS-T in the dark. Cells were then counterstained with DAPI using 1 μg/mL DAPI (Thermo Scientific, D1306) for 1 minute at room temperature. Cells were then washed 3x with 1x PBS-T. Coverslips were then mounted to the slides using ProLong Gold antifade mounting media (Thermo Scientific, P10144) and sealed with clear nail polish. Images were taken with a 60x objective on a Nikon Ti-E microscope with Andor Zyla CMOS camera using the APC and DAPI channels.

### Phase-contrast microscopy

HCI-EC-23 and Ishikawa cells were grown in regular media until roughly 80% confluent. Photomicrographs of cells were taken using a 20x objective on a Life Technologies EVOS FL microscope.

### In vivo xenograft model

All animal experiments were conducted following approved University of Utah Institutional Animal Care and Use Committee guidelines and regulations and are reported in accordance with ARRIVE guidelines. A total of 14 mice (6-9 weeks old; 17-27 g) were used in the *in vivo* xenograft growth study. Mice were housed in a temperature and humidity-controlled room with 12 hour light-dark cycles. Mice had *ad libitum* access to standard rodent food and water for the course of the study. Ovariectomized NRG (NOD.Cg-*Rag1^tm1Mom^ Il2rg^tm1Wjl^ ISzJ*, Jackson Laboratory) mice were implanted with beeswax pellets containing 0.57 mg E2 or combination of 0.57 mg E2 and 10 mg P4 under the skin on the back, between the shoulder blades. Beeswax hormone pellets were made as described previously[79]. Beeswax pellets have been shown to release hormones with a linear increase for the first 7 days post-implantation followed by sustained release with increased serum levels detected for at least 10 weeks[80]. Mice either received E2 pellets (n = 5), E2 + P4 pellets (n = 5), or no pellets (control, n = 5). Mice were then injected with 1×10^6^ HCI-EC-23 cells into the flank. 32 days after cell injection, mice were sacrificed and tumors excised and weighed. Tumors were measured using calipers starting 10 days after injection and tumor volume was calculated using the following equation: Volume = (Length × Width^2^)/2.

### Data access

Raw and processed sequencing data for RNA-seq and ChIP-seq experiments generated in this study are available under the NCBI Gene Expression Omnibus (GEO; http://www.ncbi.nlm.nih.gov/geo/) accession numbers: GSE210123 (RNA-seq) and GSE210122 (ChIP-seq). In consultation with the NCI Office of Data Sharing, only the germline-filtered MAF file describing presumed somatic mutations is available as **Supplemental File S1**. The STR cell line authentication profile for HCI-EC-23 can be provided at the time of cell line distribution or upon request.

## Supporting information

Supplemental File S1 (tab delimited)

## Acknowledgments

This research was supported by the National Institutes of Health awards R01 HG008974 to Jason Gertz and R01 HG008974S1 to Krystle Osby, the National Institutes of Health Ruth L. Kirschstein National Research Service Award from the National Human Genome Research Institute T32 HG008962 to Craig Rush, and the Huntsman Cancer Institute, including the Breast and Gynecological Cancer Center. Research reported in this publication utilized the Preclinical Research, Biorepository and Molecular Pathology, and High-Throughput Genomics Shared Resources at the Huntsman Cancer Institute at the University of Utah and was supported by the National Cancer Institute of the National Institutes of Health (P30CA042014). Research reported in this publication utilized the Cell Imaging and DNA Sequencing Core Facilities at the University of Utah. We thank the patients who volunteered their tumor specimens for this project, along with the physicians and medical staff involved in their treatment. We thank members of the Gertz lab for their insightful comments and suggestions.

## Notes

### Competing Interest Statement

The authors have declared no competing interest.

### Summary of Updates

The revised manuscript contains additional data and analysis looking at H3k27ac and ER genomic binding patterns. Other minor edits were made to improve clarity.

