## Supplemental Figures for "Characterization of a novel estrogen- and progesterone-responsive endometrial cancer cell line: HCI-EC-23"

[1]Department of Oncological Sciences, Huntsman Cancer Institute, University of Utah, Salt Lake City, UT, USA
[2]Preclinical Research Resource, Huntsman Cancer Institute, University of Utah, Salt Lake City, UT, USA
[3]Department of Biochemistry and Molecular Biology, University of Kansas Medical Center, Kansas City, KS, USA
[4]Division of Molecular Medicine, Departments of Internal Medicine and Obstetrics and Gynecology, University of New Mexico Comprehensive Cancer Center, University of New Mexico Health Sciences Center, Albuquerque, NM, USA
[5]Department of Biochemistry and Molecular Biology, Holden Comprehensive Cancer Center, Carver College of Medicine, University of Iowa, Iowa City, IA, USA
[6]Department of Obstetrics and Gynecology, University of Iowa, Iowa City, IA, USA

Additional Supplemental Files:
Supplemental Table S1. Differentially expressed genes.
Supplemental File S1. Mutation annotation file (MAF) of HCI-EC-23 filtered variants.

### Supplemental Figure S1

**a**

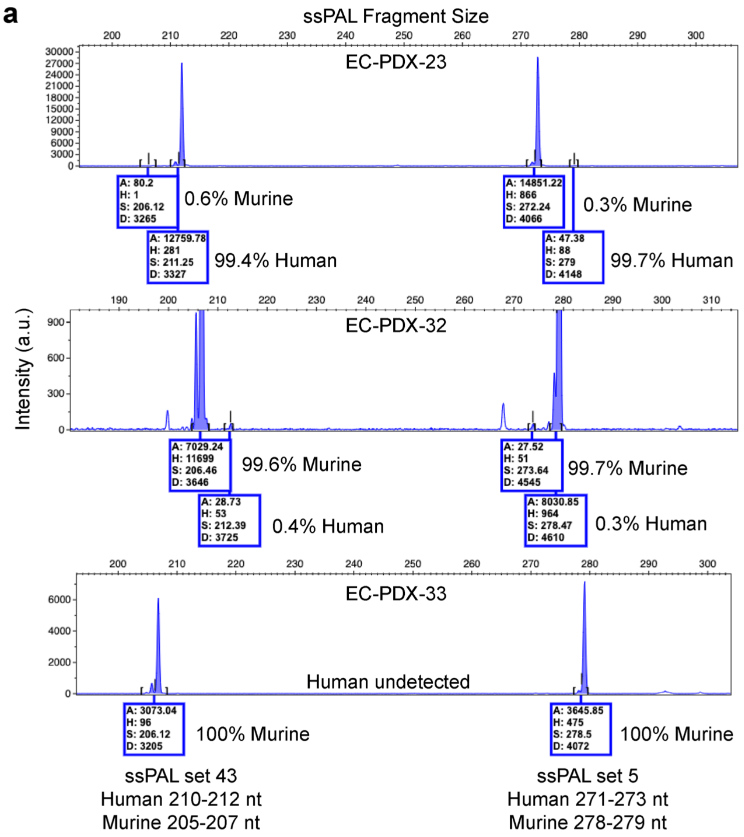


**b**

| IDEXX Species-specific PCR Evaluation | Result |
|---|---|
| mouse | Negative |
| rat | Negative |
| human | Positive |
| Chinese hamster | Negative |
| African green monkey | Negative |

| IDEXX Mycoplasma Evaluation (PCR) | Result |
|---|---|
| *Mycoplasma sp.* | Negative |

**c**

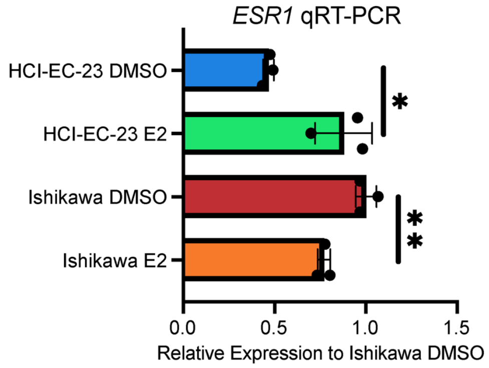


**Supplemental Figure S1. Purity of HCI-EC-23 and phenotypic comparison to the Ishikawa cell line (related to Figure 1). (a)** Fragment traces from ssPAL of sets 43 (left) and 5 (right) for EC-PDX-23 (top), EC-PDX-32 (middle), and EC-PDX-33 (bottom). Shown are the calculated areas for each murine and human amplicon along with relative percentage. **(b)** Results of commercial testing on HCI-EC-23 by IDEXX for various species (top) and mycoplasma (bottom) contamination. **(c)** Bar plot showing *ESR1* expression as assayed by qRT-PCR in DMSO- or 10 nM E2-treated HCI-EC-23 and Ishikawa cell lines. Data were normalized to internal control *CTCF* and then normalized to DMSO-treated Ishikawa. Error bars represent standard deviation. *p<0.05, **p<0.01.

### Supplemental Figure S2

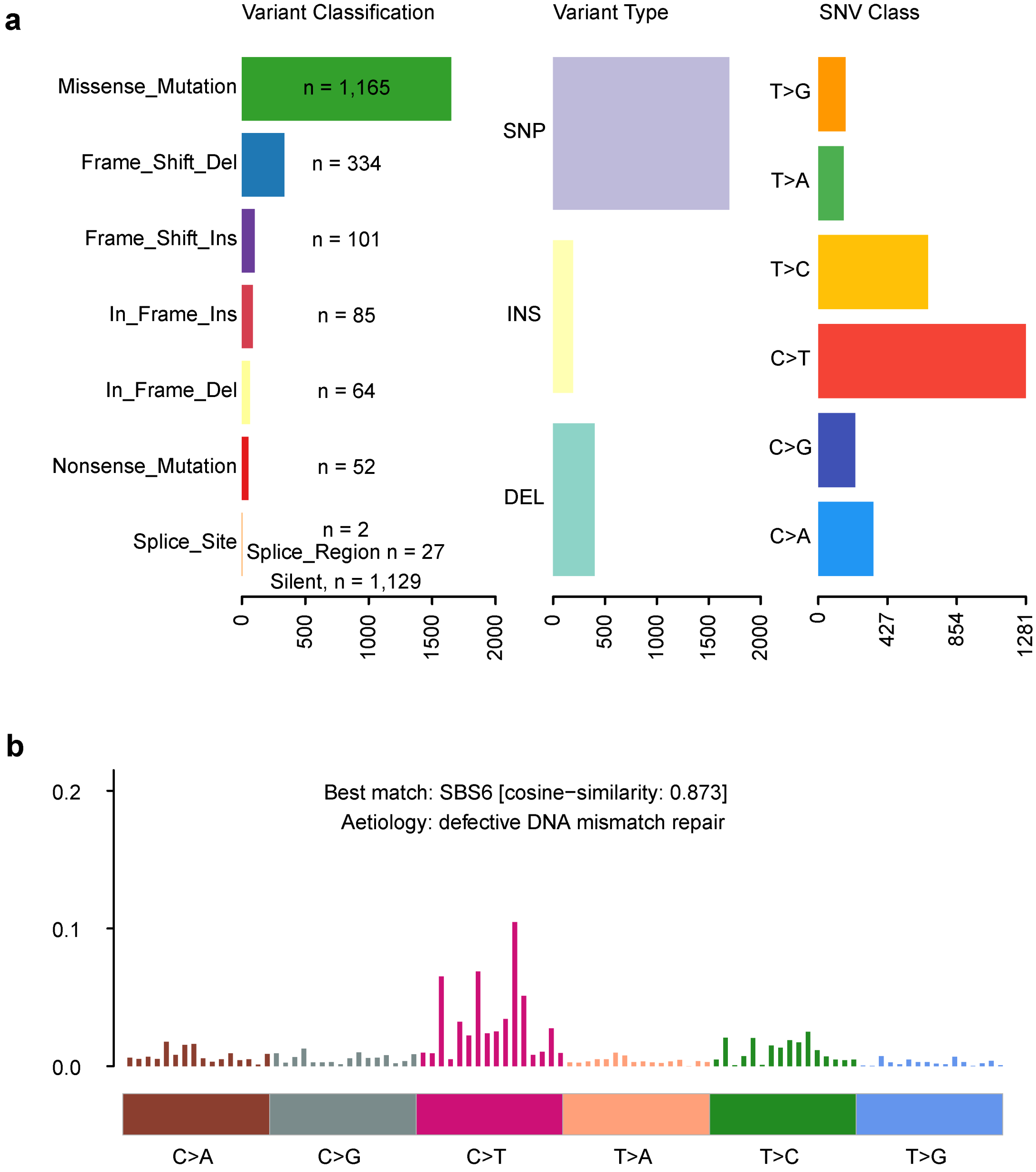


**Supplemental Figure S2. Variant class breakdown of HCI-EC-23 mutations and relationship to defective DNA mismatch repair (related to Figure 2). (a)** Bar charts displaying the number of variants within specific variant classifications (left), variant type (middle), and base change for SNVs (right). Variant classifications shown are missense, frameshift deletions/insertions, inframe deletions/insertions, nonsense mutations, splice site, splice region, and silent mutations. Variant types are categorized as SNP, insertion, or deletion. **(b)** Mutational signature identified in HCI-EC-23 and its match to SBS6 with a cosine-similarity of 0.873, indicative of DNA mismatch repair defects.

### Supplemental Figure S3

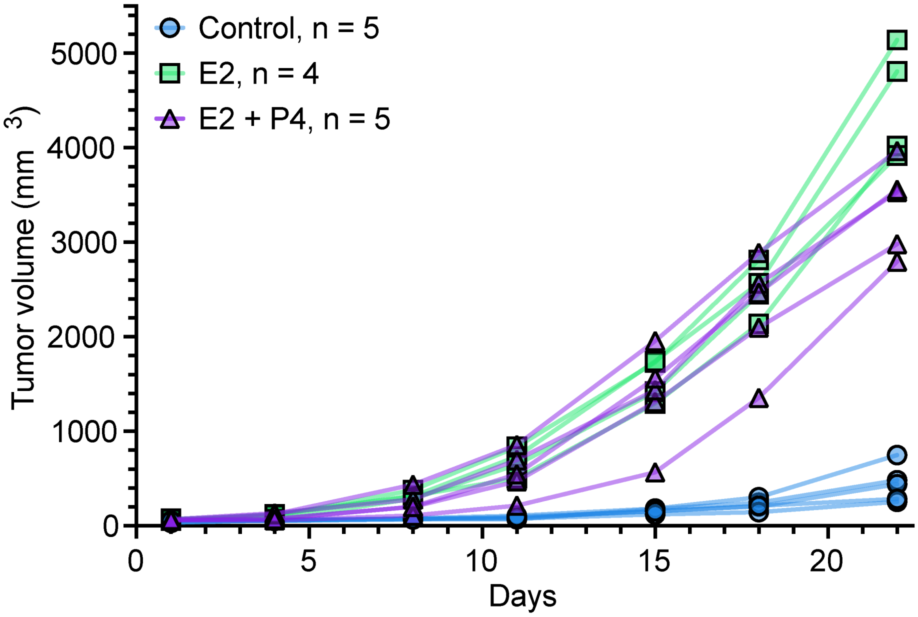


**Supplemental Figure S3. Individual xenograft tumor growth (related to Figure 3).** Line graph showing individual xenograft tumor growth of HCI-EC-23 xenografts in ovariectomized, immunocompromised mice. 1x10^6 cells were injected into the flank of mice treated with E2 (n=4), E2+P4 (n=5), or control (n=5) pellets. Day 1 represents the first day of measurement and was 10 days post-injection.

**Supplemental Figure S4**

**a**

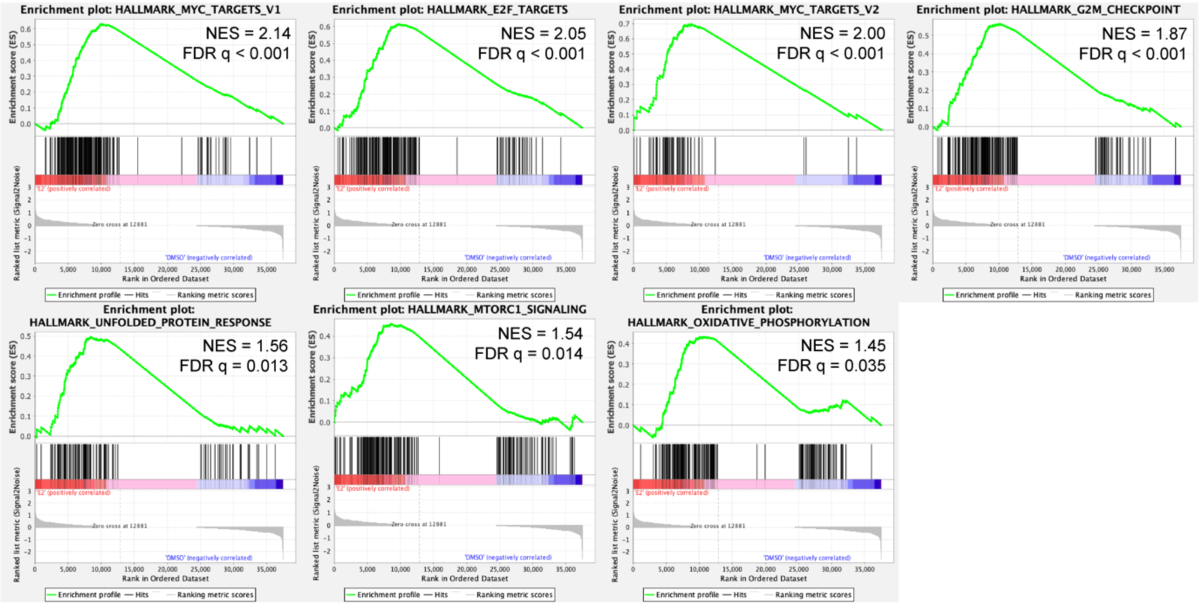


**b**

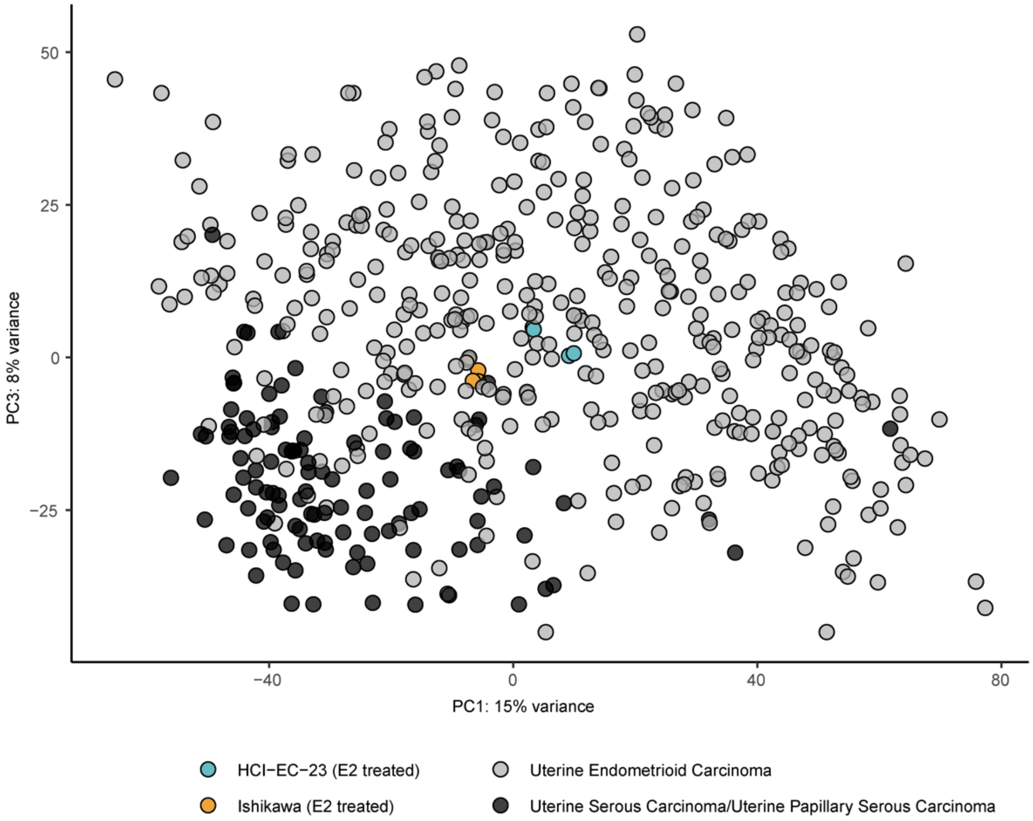


**Supplemental Figure S4. Additional enriched pathways in HCI-EC-23 E2-treated RNA-seq differentially expressed genes and comparison of HCI-EC-23 and Ishikawa RNA-seq to TCGA UCEC cohort (related to Figure 4). (a)** Gene set enrichment analysis (GSEA) plots for additional hallmark GSEA sets shown in Figure 1B. NES is the normalized enrichment score. FDR q-value is the false discovery rate. **(b)** Principal component analysis (PCA) plot of RNA-seq data for E2 treated HCI-EC-23 (blue), E2 treated Ishikawa (orange), TCGA uterine endometrioid carcinoma (gray), and TCGA uterine serous/papillary serous carcinoma. The serous and papillary serous tumors segregate from the majority of endometrioid tumors. The HCI-EC-23 and Ishikawa cell lines cluster with the endometrioid tumors.

**Supplemental Figure S5**

**a**

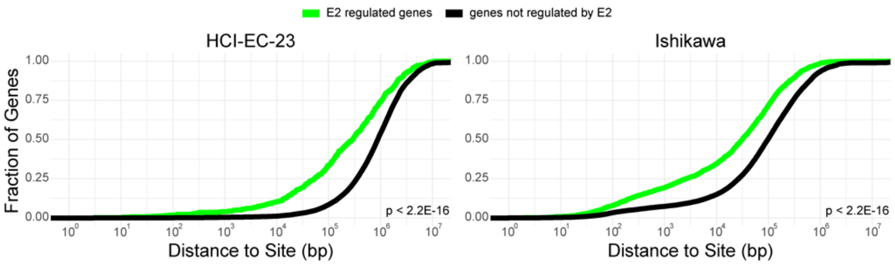


**b**

Shared ER ChIP-seq Sites

| Motif Found | Known or Similar Motif | HCI-EC-23 Estrogen Receptor ChIP E-Value Replicate 1 | HCI-EC-23 Estrogen Receptor ChIP E-Value Replicate 2 | Ishikawa Estrogen Receptor ChIP E-Value Replicate 1 | Ishikawa Estrogen Receptor ChIP E-Value Replicate 2 |
|---|---|---|---|---|---|
| 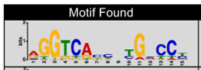 | ERE (ER) | 4.8E-691 | 3.0E-620 | 8.6E-572 | 3.0E-562 |
| 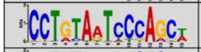 | PITX2 | 5.5E-25 | 1.1E-18 | 4.9E-36 | n.s. |
| 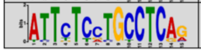 | ZN770, MAF, ZSC22, SMAD3 | 1.4E-14 | 1.1E-14 | 1.8E-17 | 2.1E-06 |

HCI-EC-23-Specific ER ChIP-seq Sites

| Motif Found | Known or Similar Motif | E-Value Replicate 1 | E-Value Replicate 2 |
|---|---|---|---|
| 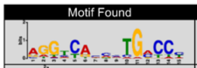 | ERE (ER) | 1.2E-242 | 1.8E-224 |
| 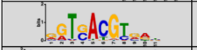 | ATF3 | 2.8E-11 | 1.2E-15 |
| 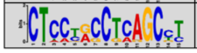 | ZN770, ZSC22, ZFX, CRX | 5.5E-07 | 4.1E-07 |

Ishikawa-Specific ER ChIP-seq Sites

| Motif Found | Known or Similar Motif | E-Value Replicate 1 | E-Value Replicate 2 |
|---|---|---|---|
| 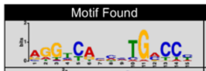 | ERE (ER) | 4.9E-1022 | 2.1E-905 |
| 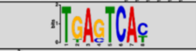 | AP-1 | 5.1E-165 | 6.4E76 |
| 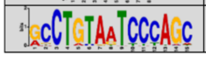 | PITX2, ZN264 | 3.1E-103 | 5.8E-60 |

**c**

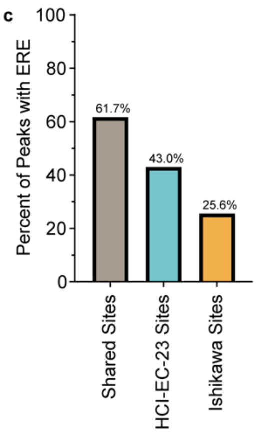


**d**

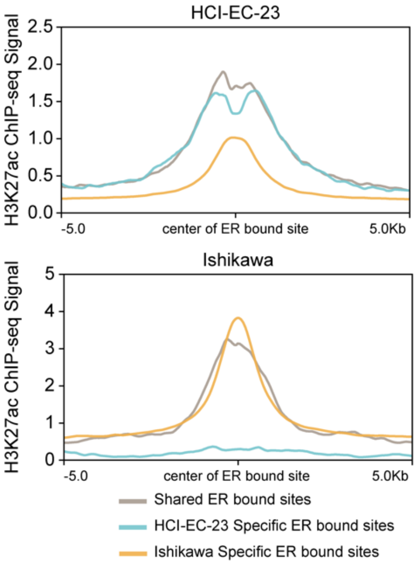


**e**

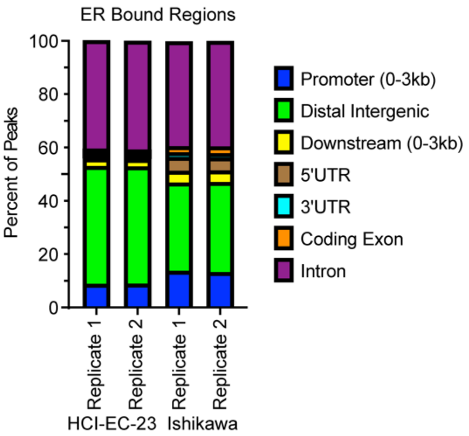


**Supplemental Figure S5. Motif analysis and genomic distribution of ER ChIP-seq peaks in HCI-EC-23 and Ishikawa (related to Figure 5). (a)** Cumulative distance plots show the fraction of E2 regulated and not regulated genes with a ER bound site within specified distance from their transcription start sites. Shown are distances for E2 regulated and not regulated genes to ER bound sites in HCI-EC-23 (left) or Ishikawa (right). Wilcoxon signed rank test p value is shown. **(b)** Top 3 motifs identified across replicates in shared (top), HCI-EC-23-specific (middle), and Ishikawa-specific (bottom) ER ChIP-seq peaks. Shown are the identified motif logo along with factors with known or similar motifs. E-values for statistical strength of motif within each set is shown by replicate. **(c)** Bar plot showing percentage of peaks harboring an ERE motif for shared (left), HCI-EC-23-specific (middle), and Ishikawa-specific (right) ChIP-seq peaks. **(d)** Profile plots are shown for H3K27ac ChIP-seq signal centered at ER bound sites. Profiles are colored according to category of ER bound site either shared, HCI-EC-23 specific, or Ishikawa specific, as indicated in the legend. Plots represent averaged signal between two ChIP-seq replicates. **(e)** Bar plots show genomic distribution of ER ChIP-seq peaks. Shown are each replicate for HCI-EC-23 and Ishikawa.

**Source Data Figure 1a**

Raw 800CW channel (Santa Cruz α-ERα sc-543)

70 kD
50 kD
38 kD

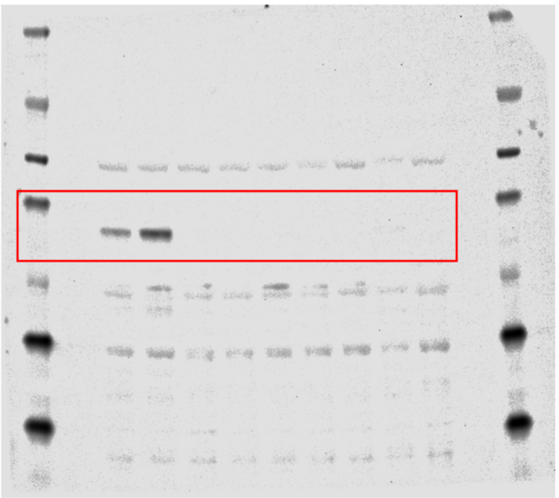

Raw 680RD channel (Santa Cruz α-Actin sc-47778)

70 kD
50 kD
38 kD

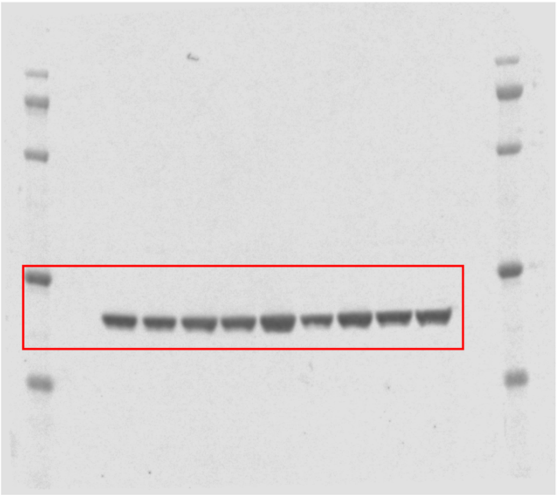

**Source Data for Figure 1a.** Provided are the raw blot images for the western blots in Figure 1a. Red boxes indicate cropped regions used for Figure 1a.
